# Mapping the human praxis network: an investigation of white matter disconnection in apraxia

**DOI:** 10.1101/2020.04.14.041442

**Authors:** Hannah Rosenzopf, Daniel Wiesen, Alexandra Basilakos, Grigori Yourganov, Leonardo Bonilha, Christopher Rorden, Julius Fridriksson, Hans-Otto Karnath, Christoph Sperber

## Abstract

Left hemispheric cerebral stroke can cause apraxia, a motor-cognitive disorder characterised by deficits of higher-order motor skills such as the failure to accurately produce meaningful gestures. This disorder provides unique insights into the anatomical and cognitive architecture of the human praxis system. The present study aimed to map the structural brain network that is damaged in apraxia. We assessed the ability to perform meaningful gestures with the hand in 101 patients with chronic left hemisphere stroke. Structural white matter fibre damage was directly assessed by diffusion tensor imaging and fractional anisotropy mapping. We used multivariate topographical inference on tract-based fractional anisotropy topographies to identify white matter disconnection associated with apraxia. We found relevant pathological white matter alterations in a densely connected fronto-temporo-parietal network of short and long association fibres. Hence, the findings suggest that heterogeneous topographical results in previous lesion mapping studies might not only result from differences in study design, but also from the general methodological limitations of univariate topographical mapping in uncovering the structural praxis network. A striking role of middle and superior temporal lobe disconnection, including temporo-temporal short association fibres, was found, suggesting strong involvement of the temporal lobe in the praxis network. Further, the results stressed the importance of subcortical disconnections for the emergence of apractic symptoms. Our study provides a fine-grain view into the structural connectivity of the human praxis network and suggests a potential value of disconnection measures in the clinical prediction of behavioural post-stroke outcome.

## Introduction

Stroke to the left brain hemisphere often causes disorders of higher-order motor control. These disorders are summarised under the term ‘apraxia’ and include symptoms such as the inability to perform, imitate, or recognize different kinds of gestures. In the present study, we investigated apraxia in the production of meaningful gestures, including the pantomime of tool use^1,2^ and communicative gestures like ‘waving goodbye’^3,4^. For simplicity, we hereafter refer to these deficits, which belong to symptoms of the classical category of ideomotor apraxia^5^, as ‘apraxia’. Many previous studies aimed to identify the neural correlates of apraxia ^6–15^. Damage in parietal, temporal, and frontal cortical regions mainly in the left hemisphere was associated with apraxia, including the angular and supramarginal gyri (SMG^6,14,15^), middle temporal (MTG^8,10,11^) and superior temporal gyri (STG^6,15^) inferior frontal gyrus (IFG^9,13,15^), insula^9,11,14,15^, and premotor areas^13^. In the light of these findings, several theories postulated a brain network to underlie apraxia, and further studies, e.g. using electroencephalography^16^, fMRI^17,18^, and fibre tracking in healthy subjects^19^, support this assumption.

The idea that apraxia results from disruptions of a complex brain network is as old as its first case descriptions by Liepmann^5^. Although theories about the architecture of this network have undergone adaptations throughout the past century, the notion of a fronto- (temporo-)parietal praxis network still dominates the field^5,14,20–23^. However, no single, widely accepted view of how the proposed network is structured exists. Further, most network theories of apraxia so far focused on cortical nodes of hypothesized networks, while white matter contributions were a minor topic in only a few previous lesion-behaviour mapping studies. White matter tracts associated with apraxia of pantomime were found below the precentral gyrus and in ventral fibres of the extreme capsule^11,19^. A multivariate lesion-behaviour mapping study found focal lesion damage to several major fibre bundles, including fronto-parietal connections, to be associated with apraxia^14^. Garcea and colleagues^15^ mapped the praxis network in reference to both structural lesion information and indirect connectome-based lesion mapping. Their study mainly implicated disconnection of parietal and ventral temporal regions, as well as disconnection of a few superior frontal regions.

However, the identification of white matter damage based on lesion-behaviour mapping bears limitations. First, univariate statistical approaches might lack statistical power to identify a full brain network^14,24^, including relevant fibre tracks. Second, topographical structural lesion-brain inference only indirectly assesses white matter contributions to symptoms via reference to brain atlases, and conclusions depend on what brain atlas is chosen to interpret results^25^. Further, a holistic understanding of such a network is difficult to achieve. While the sum of all previous lesion-behaviour mapping studies suggests a set of frontal, parietal, and temporal grey matter nodes, marked heterogeneity between topographical results exists. It is unclear to what extent the heterogeneity should be attributed to differences in sample characteristics, methodology, or apraxia assessment, and to what extent to the above-mentioned limitations of univariate statistics.

The present study aimed to directly assess white matter damage contributions to stroke patients’ apractic deficits in the production of meaningful gestures, in contrast to previous studies that only indirectly assessed white matter disconnection. We utilised multivariate spatial statistics that potentially solve the limitations of univariate statistics in identifying brain networks^26,27^ and mapped tract-based fractional anisotropy abnormalities in chronic left hemisphere stroke patients. We aimed to identify i) direct effects of lesion damage to white matter tracts, and ii) remote or long-term lesion effects on white matter integrity, as, for example, induced by Wallerian degeneration.

## Materials and methods

### Patients and behavioural assessment

We extracted data from 101 individuals who had suffered a first-time major left hemisphere ischemic stroke from an archival database at the University of South Carolina. This database contains test results from several research studies investigating aphasia and its clinical management. Patients were included in these research studies if they were between the ages of 21-80 and at least six months post-stroke. Exclusion criteria included: i) stroke involving the right hemisphere, brainstem, and/or cerebellum, ii) other neurologic illness/injury affecting the brain, iii) history of developmental speech and/or language disorder. Of note, we included participants in the present study regardless of aphasia diagnosis, and the database also includes patients with only mild or no aphasia at all. All participants signed an informed consent form that was approved by the Institutional Review Board at the University of South Carolina or the Medical University of South Carolina. All study procedures adhered to the principles set forth by the revised Declaration of Helsinki. Descriptive statistics concerning demographic, behavioural and lesion data are shown in table 1.

**Table 1.** Demographic and clinical data of all 101 patients. WAB = Western Aphasia Battery (cut-off for diagnosis of aphasic impairment < 93.8), ABA-2 = Apraxia Battery for Adults-2nd Edition.

|  | Mean | SD | Range |
| --- | --- | --- | --- |
| **Age (years)** | 59.9 | 11.0 | 30-81 |
| **Sex: f/m** | 34/67 |  |  |
| **Lesion size (cm<sup>3</sup>)** | 128,8 | 92,8 | 2,1 – 472,4 |
| **ABA-2 Apraxia Score** | 44.3 | 8.01 | 0 – 50 |
| **WAB Aphasia Score** | 60.0 | 23.4 | 16.5 – 96.4 |
| **Time post-stroke at screening (months)** | 43.2 | 46.7 | 5 - 224 |

Participants were included in the chronic stage of stroke on average 43 months after stroke onset (SD = 47 months). Aphasia was tested with the Western Aphasia Battery-Revised (WAB-R^28^). For patients with a diagnosis of aphasia (i.e., a WAB-R Aphasia Quotient <93.8), it was ensured that their understanding was sufficient for task completion, and patients with severe comprehension difficulties were excluded. Participants were included regardless of the presence of apraxia of speech and unilateral primary motor deficits.

Apraxia was assessed using the limb apraxia subscale from the Apraxia Battery for Adults-2^29^. The test contains ten gestures that are to be executed upon oral instruction (“make a fist”, “wave good-bye”, “snap your fingers”, “throw a ball”, “hide your eyes”, “make a hitchhiking sign”, “make a pointing sign”, “salute”, “play the piano”, “scratch”; for gestures involving objects, those objects weren’t present, and participants had to pantomime their use). The gesture was executed using the dominant hand post-stroke and rated between 0 (inability to complete the gesture even after demonstration of the gesture by the examiner) and 5 for best performance (i.e., the participant provides an accurate and prompt gestural response). Four points were given if the examinee produced an incorrect gesture, but self-corrected it; three points were given if the gesture was basically correct but crude and defective; two points were given if the examinee produced the correct gesture after an additional demonstration by the examiner; one point was given if the gesture after the additional demonstration was basically correct, but crude and defective^29^. Testing was conducted by an American Speech-Language-Hearing Association (ASHA)-certified speech-language pathologist, who administered and scored all assessments.

### Imaging and image processing

Brain imaging was acquired using a 3-T Siemens Trio scanner with a 12-channel head coil. T1-weighted scans were obtained with an MP-RAGE (TFE) sequence with voxel size□=□1□mm^3^, FOV□=□256□×□256□mm, 192 sagittal slices, 9-degree flip angle, TR□=□2250□ms, TI□=□925□ms, and TE□=□4.15□ms, GRAPPA□=□2, 80 reference lines. T2-weighted scans were obtained with 3D SPACE protocol with voxel size□=□1□mm^3^, FOV□=□256□×□256□mm, 160 sagittal slices, variable flip angle, TR□=□3200□ms, TE□=□352□ms, no slice acceleration, and the same slice centre and angulation as in the T1 imaging. Fractional anisotropy (FA) was computed using diffusion tensor imaging (DTI). We acquired diffusion imaging by a mono-polar sequence with 82 isotropic (2.3mm) volumes (×10 B = 0,×72 B = 1000), TR = 4987 msec, TE = 79.2 msec, 90 × 90 matrix, with parallel imaging GRAPPA = 2, and 50 contiguous slices. The sequence was carried out in two series (41 volumes in each series) with opposite phase encoding polarity. We denoised images using MRTrix (https://www.mrtrix.org/) and undistorted them using FSL’s TOPUP and Eddy tools. We converted 4D DTI images, bvecs, and bvals to fractional anisotropy maps with FSL’s DTIFIT tool. We further processed fractional anisotropy data according to the standard procedures for tract-based spatial statistics (TBSS^30^) in FSL, with the addition that during nonlinear registration each patient’s lesion was masked to avoid distortions. TBSS projects each subject’s FA data onto a mean FA tract skeleton (Figure 1).

**Figure 1.**
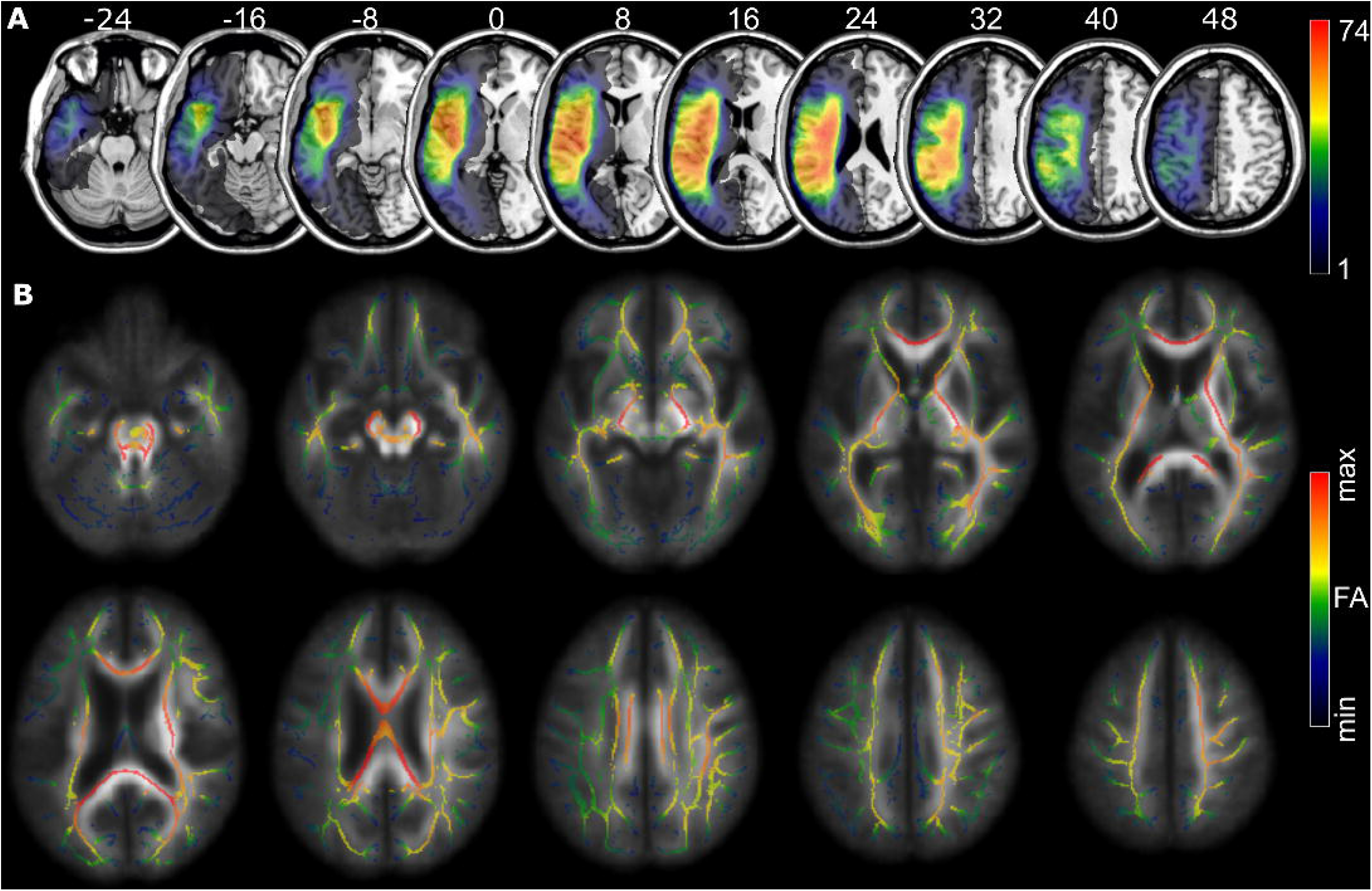
Lesion topography and mean fractional anisotropy for all 101 patients. (A) Lesion topography after enantiomorphic normalisation in SPM indicating the number of overlapping lesions per voxel. (B) Average skeletonised FA topography after normalisation and skeletonization with TBSS projected on the average non-skeletonised, normalised FA maps. Numbers above slices indicate the Z-coordinate in MNI space.

We mapped brain lesions using T2-weighted imaging. Trained experts manually delineated lesions using MRIcron (http://www.mricron.com/). We co-registered T2-weighted scans and lesion masks to T1-weighted imaging. Lesion masks were normalised by warping T1-weighted images to age-specific normalisation templates of elderly healthy controls using SPM 12 (http://www.fil.ion.ucl.ac.uk/spm/software/spm12/) and Clinical Toolbox^31^ with enantiomorphic-segmentation normalisation (Figure 1).

### Support-vector regression-based fractional anisotropy mapping

We investigated the relation between voxel-wise fractional anisotropy and apraxia scores with a multivariate topographical analysis based on support vector regression. This multivariate topographical inference method was adopted from the field of lesion behaviour mapping, where support vector regression-based lesion-symptom mapping (SVR-LSM^32^) was previously used to identify the neural correlates of apraxia of pantomime^14^. This method involved two major steps: first, we trained a support vector regression to predict behavioural scores based on fractional anisotropy data. Second, we statistically evaluated the contribution of individual voxels to the model with a permutation approach. In detail, we modelled apraxia scores with a support vector regression (SVR) with a linear kernel using voxel-wise, skeletonised fractional anisotropy topographies. Hyper-parameter C was optimised to maximise model fit and reproducibility of feature weights^32^ with five times five-fold cross-validation procedure in the range of C = [2^−20^, 2^−19^, … 2^0^, … 2^19^, 2^20^]. We evaluated model fit by SVR modelling of four-fifths of the data and assessed the correlation between the real and predicted scores in the last fifth of the data. To evaluate reproducibility, we computed the average correlation coefficient r of feature weights between each pair of cross-validation subsets and then the average r of all possible combinations of subsets. With the optimised C, we computed an SVR using data from all 101 patients and determined the feature weights, i.e. the contribution of each voxel-wise FA value to the model. A second step employed a permutation procedure to identify the features that provide a statistically significant contribution to this model. Following the SVR modelling steps as described above, voxel-wise feature weights were also computed 25000 times for data sets consisting of the original FA data, and a random permutation of the original behavioural data. Next, for each voxel, a statistical p-value was assigned to each feature weight in the original feature weight map by reference to the permutation-based feature weights. The required correction for multiple comparisons^24^ was carried out by false discovery rate (FDR) correction at q ≥ 0.05. Further analyses were conducted with the FDR-corrected topography, and with small clusters < 20 voxels removed.

### Region-based evaluation of structural disconnectivity

To identify region pair-wise disconnection associated with significant fractional anisotropy abnormalities, we evaluated the disconnectome underlying the TBSS-based SVR-LSM analysis. The exact procedure can be found in the additional online materials [http://dx.doi.org/10.17632/dcpst33wc7.2]. In short, we created a whole-brain tractogram with MRtrix (https://www.mrtrix.org/) using healthy controls’ data from the IIT Human Brain Atlas v.5.0 (https://www.nitrc.org/projects/iit/) representing the healthy connectome based on 84 ROIs of the Desikan-Kilkenny atlas and the whole-brain tractogram^33^. We then quantified the disconnectome underlying our statistical group-level results by extracting all tracts crossing the three-dimensional statistical SVR-FA-map.

In a second analysis, the clusters of significant voxels in the SVR-FA-mapping were referenced to a probabilistic white matter atlas based on diffusion tensor imaging^34^ to identify a possible affection of major fibre tracks. Probabilistic maps of long association, projection, and commissural fibres were binarised at p ≥ 0.4. Binary maps were overlaid with the SVR-FA-map and the overlapping volume was identified. Since such anatomical interpretation largely depends on the choice of white matter atlas to interpret results, we additionally referenced results to a probabilistic cytoarchitectonic atlas^35^ in the supplementary and make topographies publicly available in our online materials.

## Results

The hyper-parameter optimisation found C=2^−18^ to provide both a high reproducibility of r = 0.84 and a relatively high model fit of r² = 0.25. The permutation-based and FDR-controlled SVR-FA-map identified larger clusters of significant FA-behaviour relations in several left hemisphere regions, where increased FA was associated with more severe apraxia (Figure 2A). These included the largest clusters in parieto-temporal and temporal white matter, and further clusters in inferior frontal, parietal, and subcortical white matter.

**Figure 2.**
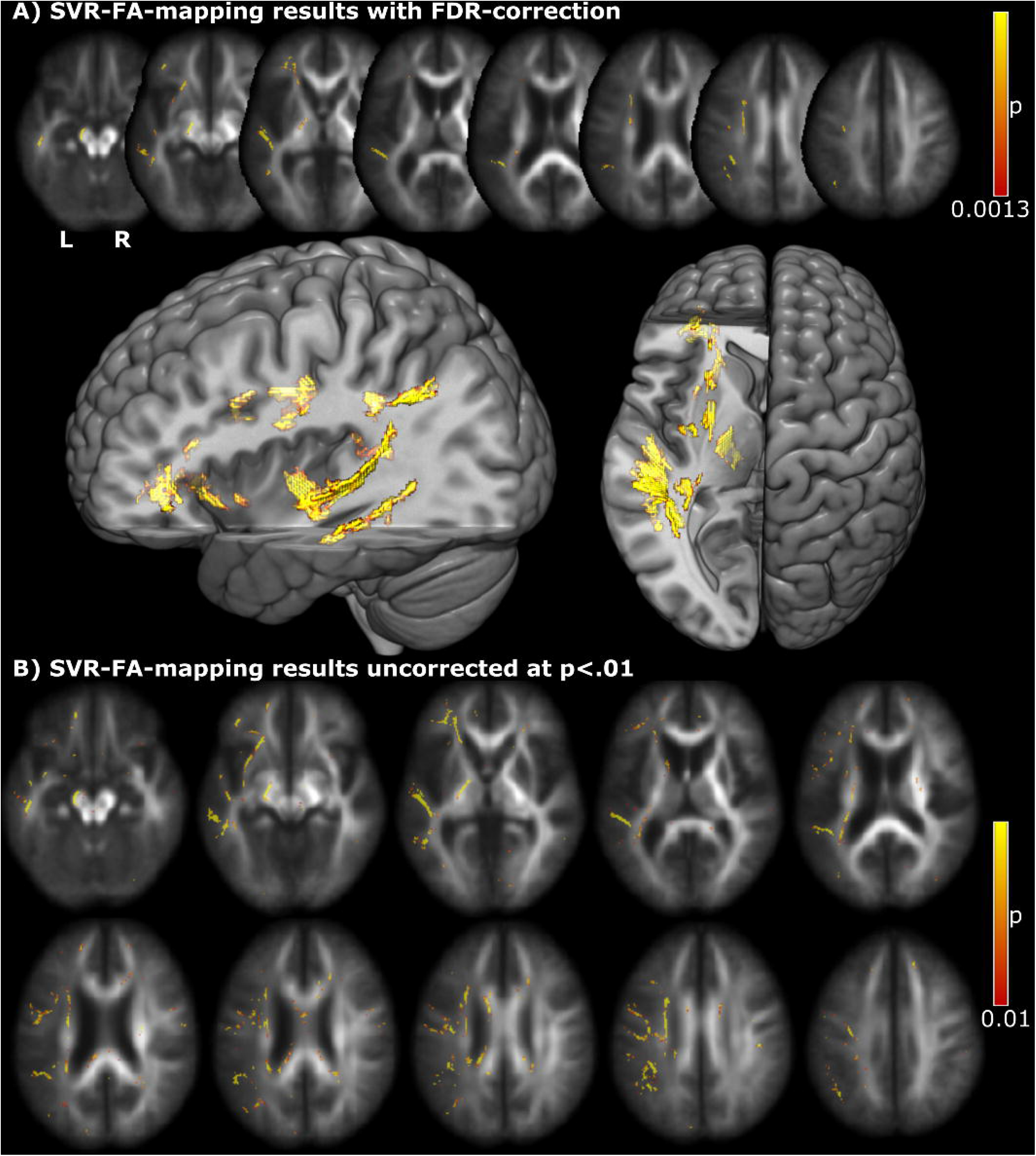
Results of the SVR-fractional anisotropy-mapping analysis. Permutation-based statistical topographies of voxels where FA values significantly contribute to apraxia are depicted (A) after false discovery rate correction with q = 0.05 (equal to p < 0.0013) and removal of small clusters < 20 voxels and (B) uncorrected at p < 0.01. 2D topographies are depicted on the un-skeletonized average FA maps; 3D topographies are depicted on the MNI152 template in MRIcroGL.

Since the present study analyzed DTI data from neurological patients and examined them for inter-regional discontinuities using DTI data from healthy subjects, we used a DTI-based atlas for referencing clusters of significant voxels to white matter anatomy (Table 2). It implicated the involvement of several major fibre tracts in apraxia, including all segments of the superior longitudinal fascicle, the uncinate fascicle, the inferior fronto-occipital fascicle, and the inferior longitudinal fascicle. Further, some significant clusters were assigned to projection fibres, including the corticospinal tract. Additionally, we referenced clusters of significant voxels to a probabilistic cytoarchitectonic atlas^35^, as documented in the online materials [http://dx.doi.org/10.17632/dcpst33wc7.2]. This analysis found a role of the inferior fronto-occipital fascicle, the corticospinal tract and callosal body.

**Table 2.** Overlap between fibre tracts as defined by Zhang *et al.*^34^ and the significant voxels found in our analysis. Abbreviations: f = fasciculus, FP = fronto-parietal, FT = fronto-temporal, PT = parieto-temporal. There was no overlap with commissural fibre tracts. Fibre tracts with less than 20mm³ overlap are not reported.

| Fibre tract category |  | Number of<br>affected voxels/<br>mm<sup>3</sup> |
| --- | --- | --- |
| Long association | Inferior fronto-occipital f. | 307 |
|  | Inferior longitudinal f. | 103 |
|  | Superior longitudinal f. (FP) | 98 |
|  | Superior longitudinal f. (FT) | 268 |
|  | Superior longitudinal f. (PT) | 474 |
|  | Uncinate fasciculus | 279 |
| Projection | Corticospinal tract | 326 |
|  | Thalamus – precentral gyrus | 38 |
|  | Thalamus – superior frontal gyrus | 127 |
|  | Thalamus – superior occipital gyrus | 36 |

In regions outside of the direct lesion areas, no significant FDR-corrected FA alterations could be detected. This means that the analysis did not identify any supra-threshold remote effects of fractional anisotropy related to apractic deficits. However, several sub-FDR-threshold clusters in the uncorrected SVR-FA-map (Figure 2B) hinted at possible remote white matter alterations. Small clusters were scattered across frontal and posterior parts of the corpus callosum and in the right hemisphere.

The region-based evaluation of structural disconnectivity identified a large number of inter-regional disconnections of varying strength (Figure 3 and Figure 4). A complete overview of disconnection metrics of all affected fibres can be found in the online materials [http://dx.doi.org/10.17632/dcpst33wc7.2]. The left hemisphere grey matter areas that displayed the highest numbers of disconnected streamlines, and that potentially execute a hub-like function in the praxis network, were found to be widely spread across the left cerebral hemisphere (see Table 3). The largest amount of disconnections was found in the basal ganglia including the putamen and caudate nucleus, in temporal areas, the inferior parietal lobe (IPL, according to the atlas parcellation^33^ consisting of inferior parietal gyrus and angular gyrus) and frontal areas. A closer look at individual pairs of regions with the highest numbers of disconnected streamlines (see Table 4, Figure 3, and Figure 4) revealed two categories of relevant connections. First, short fibres within lobes or the basal ganglia were implicated. These included several temporo-temporal, subcortico-subcortical, parieto-parietal, and fronto-frontal connections. Noteworthy, multiple connections between superior and middle temporal areas stood out. Second, long association fibres between lobes or fibres between the cortex and the basal ganglia were implicated. Among the connections with the highest amount of disconnected streamlines were temporo-parietal, fronto-subcortical, and subcortico-insular ones.

**Figure 3.**
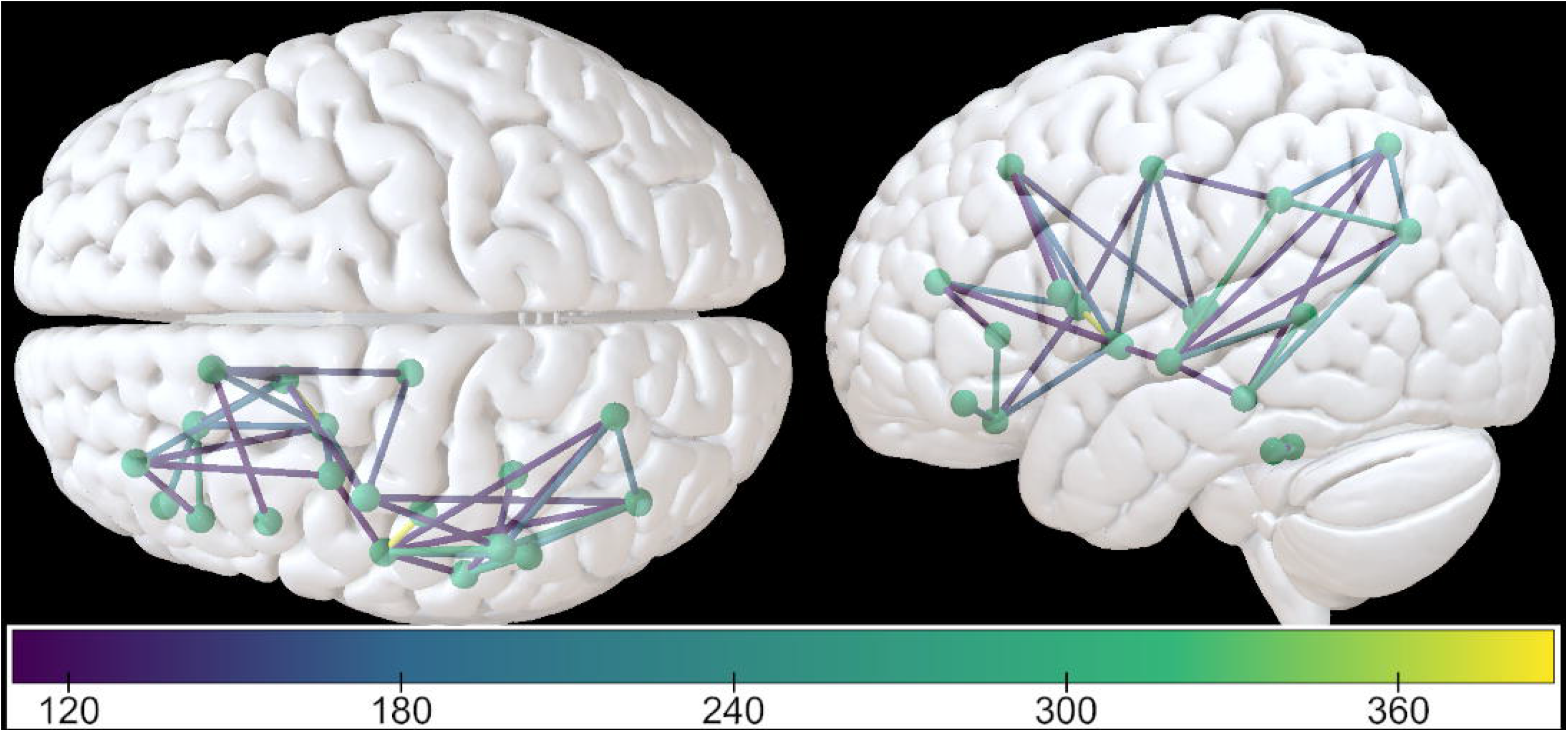
Region-based evaluation of structural disconnectivity (1). Line colour indicates the number of disconnected fibre bundles between grey matter areas in the Desikan atlas in the fibre tracking analysis in MRtrix. The threshold was set to 130 fibres to depict the strongest inter-regional disconnections. Data were visualised using Surf Ice (https://www.nitrc.org/projects/surfice/).

**Figure 4.**
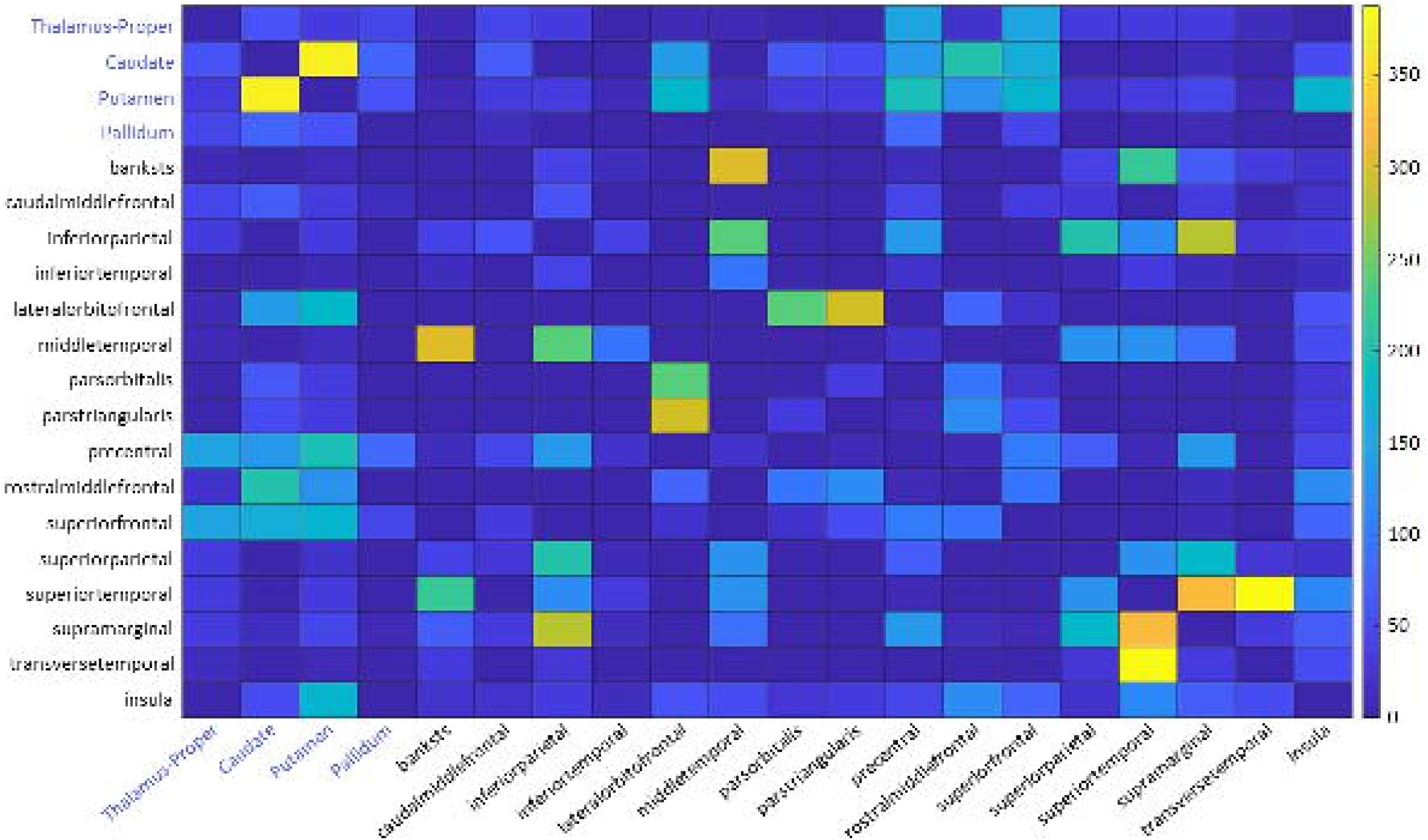
Region-based evaluation of structural disconnectivity (2). Heatmap of the 20 areas found to be most severely disconnected. Cell colour refers to the number of disconnected streamlines between the brain areas represented by the corresponding row and column. Full disconnectivity data are available in the online materials.

**Table 3.** Grey matter areas as defined by Desikan *et al*.^33^ which displayed the highest disconnection counts (> 1000) in the region-based evaluation of structural dysconnectivity.

| Brain parcellation | Number of<br>disconnections |
| --- | --- |
| Left-Putamen | 1904 |
| ctx-lh-superiortemporal | 1832 |
| Left-Caudate | 1633 |
| ctx-lh-supramarginal | 1574 |
| ctx-lh-middletemporal | 1506 |
| ctx-lh-precentral | 1487 |
| ctx-lh-inferiorparietal | 1357 |
| ctx-lh-insula | 1205 |
| ctx-lh-superiorfrontal | 1071 |
| ctx-lh-lateralorbitofrontal | 1070 |

**Table 4.** Strongest direct disconnections between grey matter pairs areas as defined by Desikan *et al.* (year) which displayed the highest disconnection counts (≥ 125) in the region-based evaluation of structural dysconnectivity.

| Disconnected brain areas | Number of disconnections |
| --- | --- |
| ctx-lh-transversetemporal - ctx-lh-superiortemporal | 388 |
| Left-Caudate - Left-Putamen | 374 |
| ctx-lh-superiortemporal - ctx-lh-supramarginal | 320 |
| ctx-lh-bankssts - ctx-lh-middletemporal | 301 |
| ctx-lh-parstriangularis - ctx-lh-lateralorbitofrontal | 292 |
| ctx-lh-supramarginal - ctx-lh-inferiorparietal | 279 |
| ctx-lh-parsorbitalis - ctx-lh-lateralorbitofrontal | 240 |
| ctx-lh-middletemporal - ctx-lh-inferiorparietal | 239 |
| ctx-lh-superiortemporal - ctx-lh-bankssts | 224 |
| ctx-lh-superiorparietal - ctx-lh-inferiorparietal | 203 |
| ctx-lh-rostralmiddlefrontal - Left-Caudate | 202 |
| ctx-lh-precentral - Left-Putamen | 195 |
| ctx-lh-supramarginal - ctx-lh-superiorparietal | 181 |
| Left-Putamen - ctx-lh-lateralorbitofrontal | 176 |
| ctx-lh-insula - Left-Putamen | 174 |
| ctx-lh-superiorfrontal - Left-Putamen | 173 |
| ctx-lh-superiorfrontal - Left-Caudate | 158 |
| ctx-lh-precentral - Left-Thalamus-Proper | 148 |
| ctx-lh-superiorfrontal - Left-Thalamus-Proper | 147 |
| ctx-lh-lateralorbitofrontal - Left-Caudate | 135 |
| ctx-lh-precentral - ctx-lh-inferiorparietal | 132 |
| ctx-lh-precentral - Left-Caudate | 131 |
| ctx-lh-precentral - ctx-lh-supramarginal | 130 |
| Left-Putamen - ctx-lh-rostralmiddlefrontal | 127 |
| ctx-lh-superiorparietal - ctx-lh-middletemporal | 125 |

## Discussion

The present analysis used MR diffusion imaging and tract-based, multivariate topographical inference to uncover white matter damage related to chronic apraxia in the production of meaningful gestures. We identified a fronto-parieto-temporal praxis network with a parieto-temporal focus. Considerable apraxia-related disconnection was found for short association fibres within the temporal lobe, as well as the parietal and the frontal lobe, and long association fibres connecting the frontal and temporal lobe with parietal areas. Notably, marked disconnection of middle and superior temporal regions was implicated, suggesting the temporal lobe as one of the main hubs in the praxis network. Further, the basal ganglia were prominently implicated with disconnection of the putamen and caudate, including connections with the frontal cortex and the insula. Significant remote effects of lesions on white matter integrity were not identified, but non-significant clusters in the right hemisphere and the callosal body hinted at potential effects that were not accessible with our methods.

### A structural praxis network

The associations found between reduced fractional anisotropy in white matter and apractic deficits largely correspond with some existing network models. A recent study on indirect connectivity measures by Garcea and colleagues^15^ mapped disconnections associated with hand posture errors in tool use gestures. They found fronto-temporo-parietal disconnection to underlie deficits in this task. They also found a focus on parietal and temporal areas. A meta-analysis by Niessen and colleagues^22^ on pantomime – i.e. meaningful gestures with an imaginary tool – points in the same direction. They suggested that apraxia of pantomime involves a network containing inferior parts of the parietal lobe, the frontal gyrus and temporal regions. We found disconnections involving the areas proposed by Niessen and colleagues as well as subcortical regions and succeeded in providing a more fine-grain description of connections underlying apraxia. Our results indicate the importance of temporo-parietal connections between lateral temporal and posterior parietal areas. Our parcellation also allowed more detailed conclusions concerning frontal region-wise disconnections. In the middle frontal gyrus, rostral and caudal portions showed high numbers of disconnected fibres. In the inferior frontal gyrus, in accordance with Garcea and colleagues^15^, particularly the pars orbitalis and pars triangularis were disconnected. Notably, the current study succeeded in identifying all these areas in association with apraxia in one single sample and with one single apraxia measure. Hence, we could show that differences in topographical results in previous studies might not only result from differences in study design or apraxia assessment, but also from the general methodology that was unsuited to identify a praxis network. The study’s multivariate approach might account for disconnections in most brain regions so far hypothesized to be relevant in apraxia, while univariate statistical approaches in previous studies tended to find only parts of this praxis network (see also Sperber et al^14^).

Identification of remote lesion effects on white matter integrity was not achieved by the FDR-corrected main analysis. However, the uncorrected statistical topography hints at possible effects. Sub-FDR-threshold clusters were apparent in the splenium and the genu of the corpus callosum. This is in line with rare cases of callosal apraxia, usually caused by damage to the splenial corpus callosum^35^ and findings in resting-state fMRI data^36^. Hence, future studies are required to synergize findings on the praxis network structure – including the present study – into a comprehensive network model that also accounts for interhemispheric communication.

### Grey matter nodes in the praxis network

The region-wise interpretation of our topographical results validated the importance of frontal, temporal, and parietal nodes in a single praxis network. Doing so, the present study integrates largely heterogeneous findings from previous lesion-behaviour mapping studies.

The IPL conventionally consists of both angular gyrus and SMG, but the atlas applied in the current study denotes an extra parcellation to SMG. The IPL was often seen as one the most central areas in a praxis network^5,21^. Our results support this notion. The SMG was among those areas with the highest number of disconnected streamlines, closely followed by the rest of the IPL, including the angular gyrus. Further, we found relevant short intra-parietal disconnections between neighbouring parietal areas, or even between subregions of the same structure. While the presence of short association fibres within a single cortical area might seem counterintuitive at first glance, it was shown that these intra-regional fibres can be identified and tracked with modern diffusion tensor imaging^32^. The parietal regions of the praxis network were previously assigned to process spatio-temporal aspects of acquired movements^38^. The parietal lobe seems to store motor programs involved in both tool use and the equivalent pantomimes^22,39^ and to code object positions in relation to oneself^39,40^. Other functions assigned to the parietal lobe include planning, predicting, and choosing appropriate actions^39^.

Disconnections of pars triangularis and pars orbitalis in the IFG are in accordance with accounts highlighting a major role of the IFG in apraxia of pantomime^9^. Goldenberg and colleagues^9^ argued that activations in the IFG might be linked to the selection of pantomime-specific action features from semantic memory. While they focused on apraxia of pantomime exclusively, the present study investigated pantomimes of object use (such as “play the piano”), and communicative gestures (such as “wave good-bye”). An alternative explanation, accounting for pantomimes and emblems, is that frontal activations are concerned with the gestures’ inherent symbolism. The IFG is popular for its role in language, a skill similarly depending on symbolism^41^. Its involvement is not restricted to spoken language but also applies to other forms of communication^42,43^. While IFG activations were greater for deaf signers of American Sign Language when attending to signs than when watching pantomimes, hearing non-signers showed higher activations for familiar pantomimes than for the unfamiliar sign language^42^. These results coincide with more recent work by Goldenberg^43^, which argued tool use pantomimes to be communicative gestures, rather than replications of real tool use. IFG activations and their connections might further indicate involvement of the mirror neuron system (MNS^42,43^), which engages the same neurons both when watching an action and executing it^45–47^. The activations found when watching communicative gestures in afore-mentioned studies might therefore also be involved in the execution of such gestures and lesions in said areas might impair processing a gesture’s inherent symbolism. Another crucial MNS region, the IPL^44^ also displayed a high number of disconnections in this study. Further damage was found in fronto-parietal fibre tracks including the fronto-parietal superior longitudinal fascicle.

Disconnection of the precentral gyrus already held an important role in classical models of apraxia. They are believed to process motor programmes forwarded from parietal areas^38^. Accordingly, EEG activity was found to begin in parietal areas and spread to left premotor areas in the course of movement preparation^16^. This role of premotor areas was argued to be generally applicable for complex motor tasks^22^. Finally, the insula, which was proposed to be critically involved in apraxia of pantomime^11,12^, displayed reduced connectivity. Like IPL and IFG, the insula is associated with the MNS, apparently connecting the frontoparietal MNS to the limbic system^44^.

While the basal ganglia are known to be an important component of motor control circuits^48^, research on apraxia indicates that subcortical areas play a less prominent role in gesture production^49^. Although cases of ideomotor apraxia following subcortical insults have been reported^50–52^, a reevaluation by Pramstaller and Marsden^49^ of all published cases of ideomotor apraxia after a subcortical stroke before 1994 revealed that only a very small fraction of lesions (8 out of 82) were isolated subcortical insults without white matter involvement^49^. A more recent study investigated sub-symptoms of ideomotor apraxia in greater detail and included both cortical and subcortical lesions^53^. Also here, at least 7 out of 9 subcortical lesions stretched into surrounding white matter. Taken together, the existing literature on subcortical area involvement in meaningful gesture production only provide limited evidence on the role of the basal ganglia in apraxia. The number of cases with isolated subcortical lesions is vanishingly low. It might therefore be that disconnections to rather than lesions in said regions are relevant for apraxia since the basal ganglia indubitably play an important role in the control of motor skills.

### Temporal contributions to a praxis network

So far, only a few lesion-behaviour mapping studies have implicated temporal regions^8,10,11,14,15^. Since the present study found strong temporo-parietal and temporo-temporal disconnections associated with apraxia, not only cortical lesions to the MTG seem to be associated with apraxia^5^, but also decreased connectivity to that very region.

Network theories of praxis most commonly assigned action recognition to temporal areas^54^. Recognition capabilities related to this area are not limited to actions, but also include objects^55^. White matter lesions affecting the posterior part of the MTG were further associated with impairing conceptual aspects of action knowledge^55^. Similarly, the relevance of MTG found in the present study might reflect conceptual issues impairing action knowledge and thus contribute to apraxia.

Results concerning the STS are again prompting towards a possible role of action recognition, more precisely, the perception of biological motion^56,57^. If this important foundation for recognizing actions^58^ plays a role in apraxia remains speculation, which requires future studies to be investigated. Further, the STG might be involved in evaluating the context-specific plausibility of action intentions^59, 60^. Since action understanding was not explicitly tested, it remains an enigma if and how understanding action and performing it relate to one another.

### There is more to praxis and its network than a single test can measure

‘Apraxia’ is often used as an umbrella term and refers to a multitude of deficits of higher-order motor skills, including, among others, the execution of meaningful gestures (Goldenberg, 2011). Deficits of higher-order motor skills can both behaviourally and anatomically dissociate into sub-deficits (Finkel et al., 2018). The present study focussed on the execution of meaningful gestures after verbal instruction. The gesture classes tested in this study, namely transitive and intransitive gestures, are known to dissociate in their cognitive load^61^and neural correlates^53,61^.

It is yet to be answered how different apraxia subtypes are related. Sub-types of apraxia occur together but are also known to dissociate occasionally^40^. The classification based on action form (e.g. here meaningful gestures) can even be further subdivided according to error type^53^. To conclude, given the complexity of praxis skills and apraxia, the delineation of the entire praxis network is not feasible within a single study. Whatever behavioural measure is chosen to access certain praxis skills; other aspects will not be assessed. Likewise, this study did not identify the entire praxis network, but the praxis sub-network related to the production of meaningful gestures – a skill commonly impaired in apraxia.

### Limitations

There are some limitations to the present study. First, disconnection severity in the region-wise interpretation refers to the absolute number of disconnected fibres, and thereby the overestimation of complete disconnections of originally faintly connected areas is avoided. However, this data presentation strategy does not provide information concerning the fraction of fibres still intact and doesn’t consider the amount of surviving fibres. Second, like in most studies on apraxia, many of the investigated patients also suffered from aphasia. This might create complications in disentangling lesion foci of these symptoms. However, the present patient sample was recruited independent of aphasia diagnosis, it included a wide range of clinical variance both for aphasia and apraxia, and the multivariate statistical inference approach was utilised to map the clinical variance of the apraxia score. Therefore, it can be assumed that the analysis indeed significantly reflects the anatomy of apraxia. Notably, this point reflects a general limitation of lesion-behaviour mapping^27^, which is not necessarily overcome by covariate control^62^. However, the complementary use of neuroscientific methods has the potential to alleviate this issue^63^. Third, the present sample was tested in the chronic post-stroke stage at least several months after stroke. This allowed us to also account for chronic long-term changes in structural network integrity. On the other hand, post-stroke brain plasticity might have altered common brain anatomy. The current study provides insights into the hyper-critical parts of the praxis network, whose damage cannot be easily compensated by brain reorganisation. Hence, to fully understand the human praxis network, acute neurological studies or studies on healthy subjects would synergise well with the current study.

## Conclusion

The present study suggests the involvement of frontal, temporal, and parietal cortical brain regions connected by short and long association fibres into a complex fronto-temporo-parietal network. A striking finding was the importance of temporal regions and temporo-temporal short association fibres, suggesting a possible hub-like function of the temporal lobe in praxis skills. Further our results implicated several cortico-subcortical connections, albeit subcortical structures were rarely implicated in previous lesion mapping studies. It seems that disconnections rather than lesions to subcortical areas might be relevant for the emergence of apractic symptoms. A future challenge is to deepen our understanding of all the essential cognitive functions underlying apraxia and to tidentify the parts of the fronto-temporo-parietal network that constitute their neural correlates. As a clinical implication of the present study, our findings suggest that long-term stroke outcome prediction based on imaging data might profit from topographical variables representing white matter damage.

## Data availability statement

All analysis scripts and resulting topographies are publicly available at http://dx.doi.org/10.17632/dcpst33wc7.2. The clinical datasets analysed in the current study are not publicly available due to the data protection agreement approved by the local ethics committees and signed by the participants. They are available from Julius Fridriksson on reasonable request.

## Author contributions

C.S., H.R., and H.O.K. contributed to the conception and design of the study; A.B., G.Y., L.B., C.R., J.F., H.R., C.S. and D.W. contributed to the acquisition and analysis of data. H.R., C.S. and D.W. contributed to drafting the manuscript and figures; H.O.K., D.W., A.B., G.Y., L.B., C.R., J.F critically revised the manuscript.

