## Supplementary for "Mapping the human praxis network: an investigation of white matter disconnection in apraxia"

### Supplementary material

### 1 Supplementary table Overlap topographical results with a probabilistic cytoarchitectonic atlas

Overlap between fibre tracts according to Juelich probabilistic cytoarchitectonic atlas (Bürgel *et al.* 2006) and the voxels found to be significant in our analysis. Fibre tracts with less than 20mm³ overlap are not depicted.

| Affected fibers | Number of affected voxels/mm³ |
| --- | --- |
| Acoustic radiation | 68 |
| Callosal body | 94 |
| Corticospinal tract | 779 |
| Inferior occipitofrontal fascicle | 37 |
| Optic radiation | 39 |

### 2 Overview tractogram creation

Evaluation of structural disconnection was assessed by generating a white matter link-wise disconnectome. The goal was to assess which connections between region of interest (ROI) pairs were altered due to white matter disruption found in the analysis. Grey matter ROIs were defined according to the Desikan-Killiany atlas and its 84 ROIs retrieved from the IIT Human Brain Atlas (v.5.0) (https://www.nitrc.org/projects/iit/ ; [Zhang & Arfanakis, 2018]). Disconnections were quantified for the statistical map resulting from the SVR-Fa-mapping analysis shown in Figure 2. The creation of the tractogram and the connectome were conducted in MRtrix3 (https://www.mrtrix.org/; Tournier *et al.*, 2012). We used the Spherical harmonic (SH) coefficients of the IIT_HARDI.nii template file (Varentsova *et al.*, 2014) from the IIT Human Brain Atlas (v 5.0.) as input to perform tractography. To prepare the mask for streamline seeding we started with the command *5tt2gmwmi* on the

IIT_fornix_fixed_5tt_file_for_ACT_tractography.nii, available from the IIT Human Brain Atlas (v.5.0). The mentioned template contains a 5 tissue type segmented anatomical image, which enabled us to anatomically constrain the streamlines and hence, to improve the accuracy of streamline termination (Smith *et al.*, 2012). The *tckgen* command was used to generate streamlines. To do so, we used the default probabilistic algorithm (*iFOD2*, Tournier *et al.*, 2010) with anatomical constrained deconvolution. 10 million streamlines were seeded at the grey matter – white matter boundary. Additionally, *backtrack* (Smith *et al*., 2012) allowed to resample streamlines that had been rejected previously, in case of a poor structural termination. Next, *tcksift* was used. The command executes filtering of tracks with the so-called spherical-deconvolution informed algorithm (Smith *et al.*, 2013). This transforms the streamline densities to match the FOD lobe integrals and reduces overestimation of longer tracks. Secondly, it turns the number of streamlines between two regions into a proportional estimate of the cross-sectional area of fibres that connect the two concerned regions. Consecutively, the connectome of the resulting whole-brain tractogram was created with *tck2connectome*, by quantifying the number of streamlines between any two Desikan-Kiliany ROIs (Desikan *et al.*, 2006). Additionally, we created a second connectome file for our SVR-FA-mapping statistical result by removing first all streamlines of the whole-brain tractogram running through the statistical map with *tckedit*. Then, we again quantified the number of streamlines in a same way as for the 'healthy' connectome and took the difference between both. This allows us to specifically quantify which areas of the brain are structurally disconnected or altered (i.e. the disconnectome) with reference to our statistical findings.
